# miR-424(322)^∼^503 impairs colon cancer progression driven by PTEN deficiency

**DOI:** 10.64898/2025.11.28.691183

**Authors:** Maria Vidal-Sabanés, Núria Bonifaci, Raúl Navaridas, Joaquim Egea, Mario Encinas, Ruth Rodriguez-Barrueco, Jose M. Silva, Xavier Matias-Guiu, Jordi Tarragona, David Llobet-Navas, Xavier Dolcet

## Abstract

Colorectal cancer (CRC) is a leading cause of cancer-related morbidity and mortality worldwide, with molecular subtypes and signaling pathways playing critical roles in its progression. The miR-424(322)^∼^503 cluster, comprising miR-424 and miR-503, has been implicated in various malignancies, exhibiting dual roles as tumor suppressors or oncogenes depending on the context. However, its function in CRC remains poorly understood. This study investigates the role of the miR-424(322)^∼^503 cluster in CRC driven by PTEN deficiency using genetically modified mouse models. Our findings reveal that the loss of miR-424(322)^∼^503 significantly exacerbates CRC progression in PTEN-deficient mice. Double knockout (dKO) mice lacking both PTEN and miR-424(322)^∼^503 exhibited a higher number and larger size of colorectal lesions compared to PTEN-deficient counterparts. Histological analysis demonstrated increased severity of dysplasia and adenocarcinoma development in dKO mice. Mechanistically, while Wnt/β-catenin signaling remained unaltered, transcriptomic analyses highlighted dysregulation of MAPK and TGFβ pathways, alongside epithelial-to-mesenchymal transition (EMT)-related gene signatures. Protein-level validation confirmed hyperactivation of MAPK (ERK1/2 and p38) and TGFβ signaling, as well as elevated cyclin D1 expression in dKO colonic tissues. These results underscore the tumor-suppressive role of the miR-424(322)^∼^503 cluster in CRC by modulating key oncogenic pathways such as MAPK and TGFβ. Our study provides novel insights into the interplay between PTEN loss and miRNA regulation in CRC pathogenesis.

## INTRODUCTON

Worldwide, colorectal cancer (CRC) is the fourth most common type of cancer and the third most deadly, taking into account men and women. In fact, in 2022, 1,926,425 new cases of colorectal cancer were diagnosed and 904,019 deaths were recorded ^1,2^. Histologically, the vast majority of colorectal cancers are adenocarcinomas originating in the epithelial cells of the mucosa of the colon or rectum. However, in the histological classification established in the fifth edition of the World Health Organization’s tumor classification in 2019 (WHO, 2019), there are nine more variants of this type of tumor ^3,4^. Due to the need to improve the prognostic classification provided by tumour histology, in 2015 a consensus of molecular subtypes (MSCs) was reached based on the transcriptomic analysis of a large number of primary tumours worldwide. The results obtained made it possible to study aspects related to the tumor microenvironment and metabolic, genomic and epigenetic signatures, which resulted in the establishment of four molecular subgroups ^5,6^. The molecular classification of CRC is based on genetic and epigenetic characteristics, such as mutations, microsatellite or MSI instability, methylation of CpG islands (*CpG Island Methylator Phenotype*, CIMP), chromosomal instability (*Chromosomal Instability*, CIN), alterations in the number of copies (*Copy Number Alterations*, CNA) or overactivated signaling pathways ^7^. The majority of colorectal cancers, between 70% and 90%, arise from precancerous polyps or adenomas that, over a long period of time, undergo a series of transformations until neoplasia develops ^8,9^. At the base of the colon crypts there is the proliferation and differentiation of intestinal stem cells, which allows the cell turnover necessary to replace the cells that exfoliate in the intestinal lumen ^10^. In the adenoma-adenocarcinoma sequence, cells undergo alterations in DNA repair mechanisms, which gives them a certain genomic instability that encourages the formation of a dysplastic epithelium ^9^. The onset of adenoma formation coincides with the inactivating mutation of *APC*, which provides cells with proliferative self-sufficiency through sustained activation of WNT signaling. Later, mutations in *KRAS*, *SMAD4* and *TP53* provide telomere dysfunction and chromosomal instability and even invasive and metastatic capacity, resulting in the formation of invasive adenocarcinoma ^8^. Apart from traditional adenomas, there are other precursor lesions that give rise to colorectal cancer. This is the case of serrated polyps, which represent the precursor lesion of 15% of the total CRCs, after having been transformed by the sawn pathway. This neoplastic pathway is characterized by presenting the activating mutation of *BRAF* V600E, which causes constitutive activation of MAPK signaling resulting in uncontrolled cell proliferation. After the mutation in *BRAF*, serrated tumors can evolve into two different pathways: the MSI pathway, which is based on alterations in DNA repair genes and results in a phenotype with a high MSI; and the way of *TP53*, which promotes the activation of signaling pathways such as TGF-β or EMT. It is important to note that in tumors originating from the sawn pathway, an overactivation of the WNT pathway can be found, which, unlike the traditional route, is not due to the inactivation of *APC* ^9^. the inactivating mutations of *PTEN*, which are found in 10% of RCCs, also induce constitutive activation of AKT ^11^, identified as a mutation initiating the process of colorectal carcinogenesis, alterations in *PTEN* participate in the progression of this type of tumor. In this sense, it has been seen that the loss of expression of *PTEN* is highly related to genetically unstable tumors ^12^, as well as with tumors of greater stage and with the presence of nodular metastases ^13^. Mutations in *PTEN*, moreover, often coexist with alterations in *BRAF* and, therefore, with the serrated pathway of colorectal pathogenesis that is characterized by having a high state of MSI and CIMP ^14^.

MicroRNAs (miRNAs) are small non-coding RNA molecules that regulate gene expression and play critical roles in cancer biology. The miR-424(322)^∼^503 cluster expresses miR-424(322 in mouse) and miR-503, two miRNAs belonging to the miR-16 family ^15^. Among the growing number of miRNAs implicated in cancer, miR-424(322)^∼^503 has garnered attention due to its dual role in tumorigenesis, both as a tumor suppressor and as an oncogene, depending on the cellular context. ^16–18^. This miRNA-cluster is involved in the regulation of cancer-related cellular processes such as proliferation, differentiation cellular plasticity or apoptosis ^19^ and has been found either up-or-down-regulated in many types of cancers ^16–18^.

The role of miR-424(322)^∼^503 in CRC has not been deeply explored, and as in other types of malignancies, a dual role has been reported. As oncogenic functions, it can promote proliferation and metastasis ^20,21^. As tumor suppressive miRNA-424 has been shown to reduce tumor progression^22^, angiogenesis^23^ and Epithelial-to-Mesenchymal Transition (EMT) ^24^. To date, there are no studies investigating the *in vivo* function of the miR-424(322)^∼^503 cluster in CRC using genetically modified mouse models. Here, we aimed to investigate the role of the miR-424(322)^∼^503 cluster in PTEN-loss-driven CRC.

## RESULTS

### Lack of miRNA-322^∼^503 enhances CRC tumorigenesis in Pten-deficient mice

We have recently demonstrated that miR-322^∼^503 is required for endometrial cancer (EC) progression initiated by Pten deficiency^25^. In this previous study, we analyzed the effects of miR-322^∼^503 in EC initiated by Pten deficiency by crossing a tamoxifen-inducible conditional Pten Knock-out mice (*Cre:ER*(T)*^+/-^Pten^F/F^*) with a miR-322^∼^503 knock-out mice miRNA-322/503^-/-^ to obtain four genotype combinations: *Cre:ER^(T-/-^Pten^F/F^*miRNA-322/503^+/+^ (Wt), *Cre:ER^(T/-/-^Pten^F/F^*miRNA-322/503^-/-^ (miR KO), *Cre:ER*(T)*^+/-^Pten^F/F^*miRNA-322/503^+/+^ (*Pten* KO) mice and the *Cre:ER*(T)*^+/-^Pten^F/F^*miRNA-322/503^-/-^ (*d*KO) mice (**Supplementary Figure S1**). In this previous study, we demonstrated that *d*KO mice had impaired progression of EC compared to *Pten* KO mice. While performing necropsy of mice to study the role of miR-322^∼^503 in the endometrium (**Figure 1A**), we macroscopically noticed that *d*KO mice had enlarged intestines. To examine the cause of such enlargement, intestines from mice were dissected and opened longitudinally. Macroscopic observation revealed a significant number of superficial lesions in *Pten* KO mice but, surprisingly, the number of lesions was markedly increased *d*KO mice compared to any other genotype (**Figure 1B**). We quantified the number and measured the size of macrospic lesions observed in the epithelial surface were in both the *Cre:ER*(T)*^+/-^Pten^F/F^*miRNA-322/503^+/+^ (*Pten* KO) mice and the *Cre:ER*(T)*^+/-^Pten^F/F^*miRNA-322/503^-/-^ (*d*KO) mice (**Figure 1C**). To determine if there were differences in lesion size between *Pten* KO and dKO, the volumes of each lesion were calculated. *Cre:ER*(T)*^-/-^Pten^F/F^*miRNA-322/503^+/+^ (Wt) and *Cre:ER*(T)*^-/-^Pten^F/F^*miRNA-322/503^-/-^ (*miR* KO) mice showed no lesions throughout the intestinal epithelium (**Figure 1B**). In contrast, a significant increase in the multiplicity of lesions was observed in the colons of dKO mice compared to *Pten* KO colons (**Figure 1C**). Moreover, dKO lesions exhibited a marked increase in volume compared to their *Pten* KO counterparts (**Figure 1D**). To rule out the possibility that the differences in the number of polyps were due to intestinal length, the lengths of the colons were measured and compared, showing no statistically significant differences between the two genotypes (**Figure 1E**).

**Figure 1.**
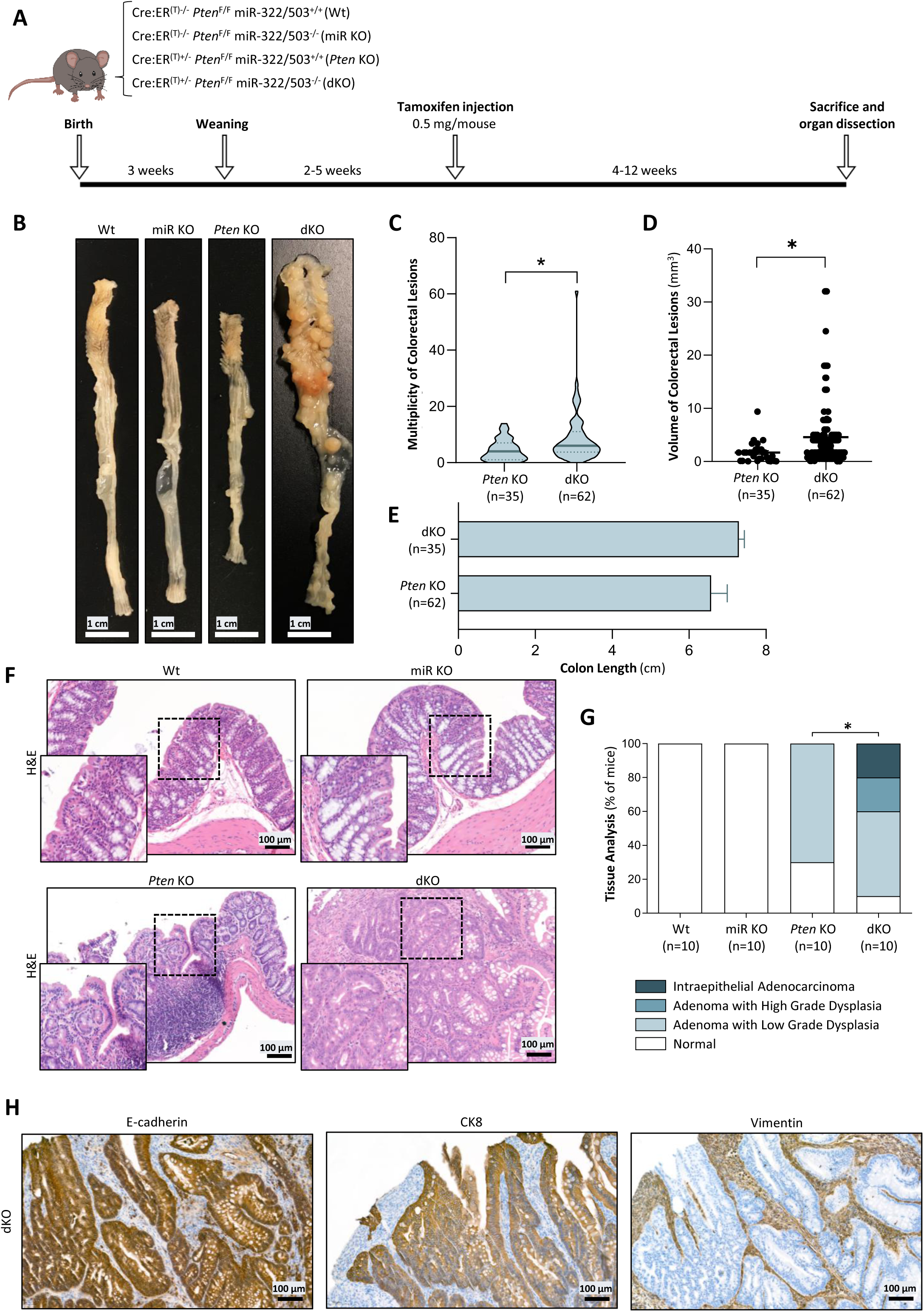
Double deficiency of *Pten* and miR-322/503 increases multiplicity, volume and complexity of colorectal polyps. **(A)** Timeline for the in vivo analysis of endometrial lesions in Cre:ER(T)^-/-^ *Pten*^F/F^ miR-322/503^+/+^ (Wt), Cre:ER(T)^-/-^ *Pten*^F/F^ miR-322/503^-/-^ (miR KO), Cre:ER(T)^+/-^ *Pten*^F/F^ miR-322/503^+/+^ (*Pten* KO) and Cre:ER(T)^+/-^*Pten*^F/F^ miR-322/503^-/-^ (dKO) mice. **(B)** Representative macroscopic images of colons from Wt, miR KO, *Pten* KO and dKO mice. Scale bar: 1 cm. **(C)** Violin plot showing multiplicity of colorectal lesions in Pten KO and dKO mice. Statistical significance was determined using t-test analysis, *p<0.05. **(D)** Volume of colorectal lesions observed in *Pten* KO and dKO mice. Statistical differences were assessed by t-test analysis, *p<0.05. **(E)** Colon length of *Pten* KO and dKO mice. Data is presented as mean ±S.E.M, according to t-test analysis. **(F)** Representative hematoxylin and eosin (H&E) staining of colorectal sections from Wt, miR KO, *Pten* KO and dKO mice. Scale bar: 100 μm (15X magnification). **(G)** Histopathological analysis of colorectal sections from Wt, miR KO, *Pten* KO and dKO mice showed in (F). Statistical analysis was determined using Chi-squared analysis, *p<0.05. **(H)** Representative immunohistochemistry images of E.cadherin, cytokeratin-8 (CK8) and Vimentin in CRC lesions od dKO mice. Scale bar: 100 μm (15X magnification).

After conducting the macroscopic analysis, the colons were embedded in paraffin for subsequent histological and immunohistochemical analysis. The histological analysis was performed on colons from Wt, *Pten* KO, *miR* KO or dKO mice. For this purpose, we carried routine hematoxylin-eosin staining to observe histopathological alterations (**Figure 1F**). Consistent with the lack of macroscopic intestinal lesions, all Wt and *miR* KO colons exhibited normal histology. On the other hand, *Pten* KO colons, 30% showed normal histology, while 70% presented adenomas with low-grade dysplasia. Finally, in dKO mice, 10% exhibited normal colorectal histology, while the remaining cases were distributed as follows: 50% with adenomas with low-grade dysplasia, 20% with adenomas with high-grade dysplasia, and 20% with intraepithelial adenocarcinomas (**Figure 1F, 1G**). IHC staining of cytokeratin, E-cadherin and vimentin, evidenced the epithelial origin of CRC developed in dKO mice (**Figure 1H**).

### Lack of Pten enhances miR-322^∼^503 expression in colorectal epithelium

In our previous study, we demonstrated that *Pten* loss caused an increase of miR-322^∼^503 expression in endometrial cells ^25^. Having found a completely antagonistic effect of miRNA deletion in the colon, we questioned whether the expression of miR-322^∼^503 in the colorectal epithelium of *Pten* KO mice was also affected. For this purpose, RNA from total epithelial tissue, RNA from polyp lesions or RNA from colon organoids was extracted from the colons of Wt or *Pten* KO mice and the miRNA expression was analyzed by quantitative PCR **(Figure 2A)**. The expression results showed an upward trend in both miRNA-322 and miRNA-503 expression in Pten KO mice compared to Wt ones (**Figure 2B**). It is important to mention that PTEN deficiency triggered upon tamoxifen injection in epithelial cells occurs in a mosaic pattern resulting in ablation only in a fraction of total epithelial cells, thereby diluting the possible effects of PTEN deficiency in miRNA expression (**Supplementary Figure S2A**). To sort this out, miRNA was extracted from colorectal polyps dissected from *Pten* KO mice, in which most cells have lost *Pten* expression (**Supplementary Figure S2B**). The results demonstrated a significant increase in the expression of both miR-322 and miR-503 in PTEN-deficient colorectal polyps (**Figure 1C**). This increase in the miRNA expression was further enhanced in colon organiods in *Pten* KO colon organoids compared to the Wt ones (**Figure 1D-E**).

**Figure 2.**
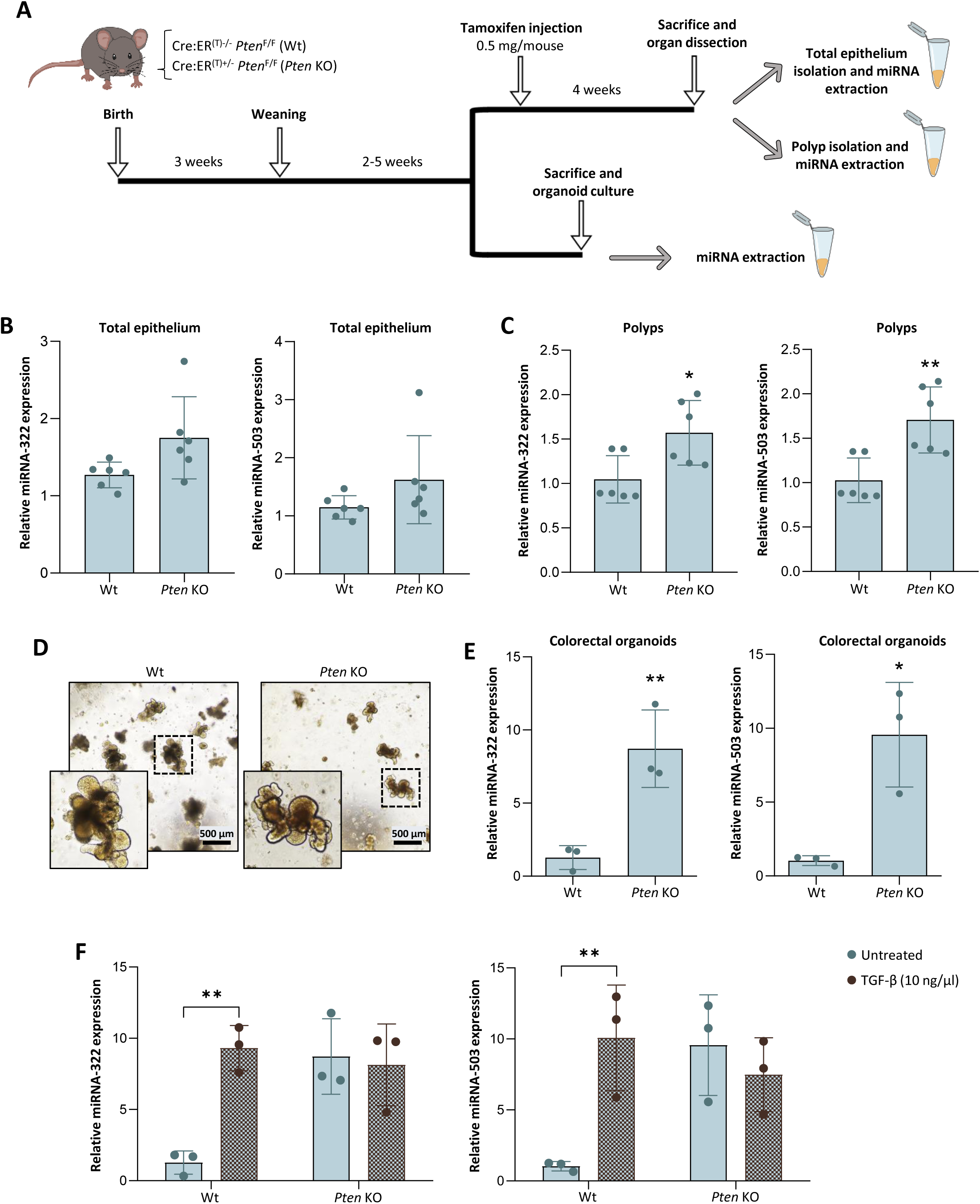
*Pten* deficiency increases miRNA-424(322)/503 cluster expression in colorectal epithelial cells. **(A)**Timeline for the analysis of miR-322 and miR-503 expression in colorectal epithelium from Cre:ER(T)^-/-^ *Pten*^F/F^ (Wt) and Cre:ER(T)^+/-^ *Pten*^F/F^ (*Pten* KO) mice. **(B)** Relative miR-322 and miR-503 expression in total colorectal epithelium or **(C)** polyps from Wt and *Pten* KO mice. **(D)** Representative images of colon organoids from Wt and *Pten* KO mice. Scale bar: 500 μm (4X magnification). **(E)** Relative expression of miR-322 and miR-503 in Wt and *Pten* KO mouse colon organoids. **(F)** Relative expression of miR-322 and miR-503 in Wt and *Pten* KO mouse colon organoids treated with 10 ng/μl TGF-β for 16 hours. Data are presented as mean ±S.E.M, according to t-test analysis, *p<0.05; **p<0.01.

In our previous study, we also demonstrated that treatment of organoids with TGFβ caused an increase of miR-322^∼^503 expression in endometrial cells ^25^. Therefore, we questioned whether this regulation was also important in colon epithelial cells. To address this question, we treated Wt and *Pten* KO colon organoid cultures with TGFβ and we measured expression of miRNA-322 and miRNA-503. As we had previously observed in the endometrium, colon organoids showed a marked increase in the expression of both miRNAs after TGFβ treatment (**Figure 1F).** These results underscore the role of PI3K/Akt and TGFβ signaling in the regulation of miR-424(322)^∼^503 cluster expression.

### CRC from double Pten and miR-322^∼^503 deficient mice do not display alterations in β-catenin signaling

It is well documented that miR-322^∼^503 plays a role in regulating the Wnt/β-catenin signaling pathway in mammary tissues through its binding to the messenger RNA of LRP6, one of the receptors of this pathway ^26^. This, combined with the critical role of this signaling pathway in colorectal cancer ^27^, leads to the hypothesis that dKO mice may suffer an hyperactivation of the Wnt/β-catenin pathway in colorectal tissue, resulting in the development of malignant lesions in this organ. To address this question, we performed immunohistochemical staining against the LRP6 receptor and β-catenin. Interestingly, no increase in either LRP6 or β-catenin was observed in dKO colons compared to *Pten* KO colons. Based on these results, the lack of miRNA-322/503 expression in PTEN-deficient colons does not alter the Wnt/β-catenin pathway, ruling it out as the cause of the increased colorectal lesions (**Figure 3**).

**Figure 3.**
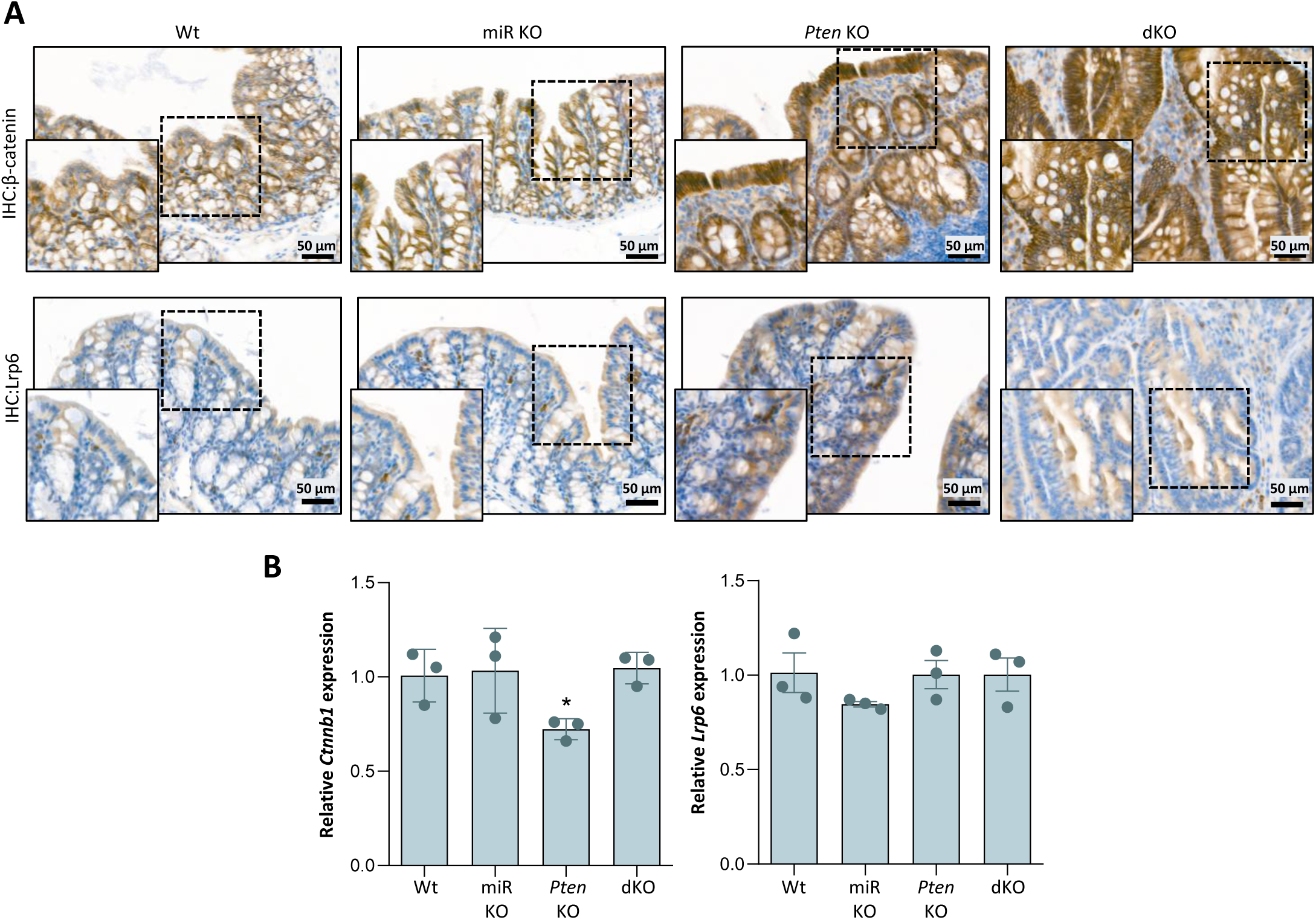
Carcinogenesis induced by lack of Pten and miRNA-322/503 is not mediated by Wnt/β-catenin signaling. **(A)** Representative immunohistochemistry images of β-catenin and Lrp6 in colon sections from Cre:ER(T)^-/-^ *Pten*^F/F^ miR-322/503^+/+^ (Wt), Cre:ER(T)^-/-^ *Pten*^F/F^ miR-322/503^-/-^ (miR KO), Cre:ER(T)^+/-^ *Pten*^F/F^ miR-322/503^+/+^ (*Pten* KO) and Cre:ER(T)^+/-^ *Pten*^F/F^ miR-322/503^-/-^ (dKO) mice. **(B)** Relative expression of *Ctnnb1* and *Lrp6* in Wt, miR KO, *Pten* KO and dKO colorectal tissue. Data are presented as mean ±S.E.M, according to t-test analysis, *p<0.05.

### Double Pten and miR-322^∼^503 deficiency regulates the expression of genes associated with MAPK signaling, TGFβ signaling and epithelial-to-mesenchymal transition

Since β-catenin did not seem to be altered in dKO mice, we pursued alternative molecular alterations that could explain colorectal carcinogenesis induced by the dual loss of miR-322^∼^503 and PTEN. To have a global view of this dual loss effects on gene expression, we performed a comprehensive mRNA of colon epithelium from Wt, *Pten* KO, *miRNA* KO or dKO mice. To investigate the transcriptional impact of miR-322^∼^503 deletion, we performed a differential gene expression analysis (DEG, log_2_ FC > abs(1.5) plus adj. FDR p< 0.05) to compare gene expression profiles across the following groups: miR KO vs. Wt, *Pten* KO vs. Wt, dKO vs. Wt, and *Pten* KO vs. dKO **(Figure 4A-B**, **Supplementary Table 1)**. As expected by the dramatic impact of double Pten and miRNA ablation in CRC development, the comparison between samples dKO and Wt exhibited the highest number of genes with significantly altered expression. However, the comparison between *Pten* KO and Wt samples also exhibited a large number of genes with significantly altered expression, indicating the profound transcriptional changes associated with *Pten* loss. Then, to find differentially regulated biological processes or molecular functions (i.e., pathway-level insights) underlying CRC development, we conducted an exploratory analysis by GSEA ^45^ using predefined gene set annotations.

**Figure 4.**
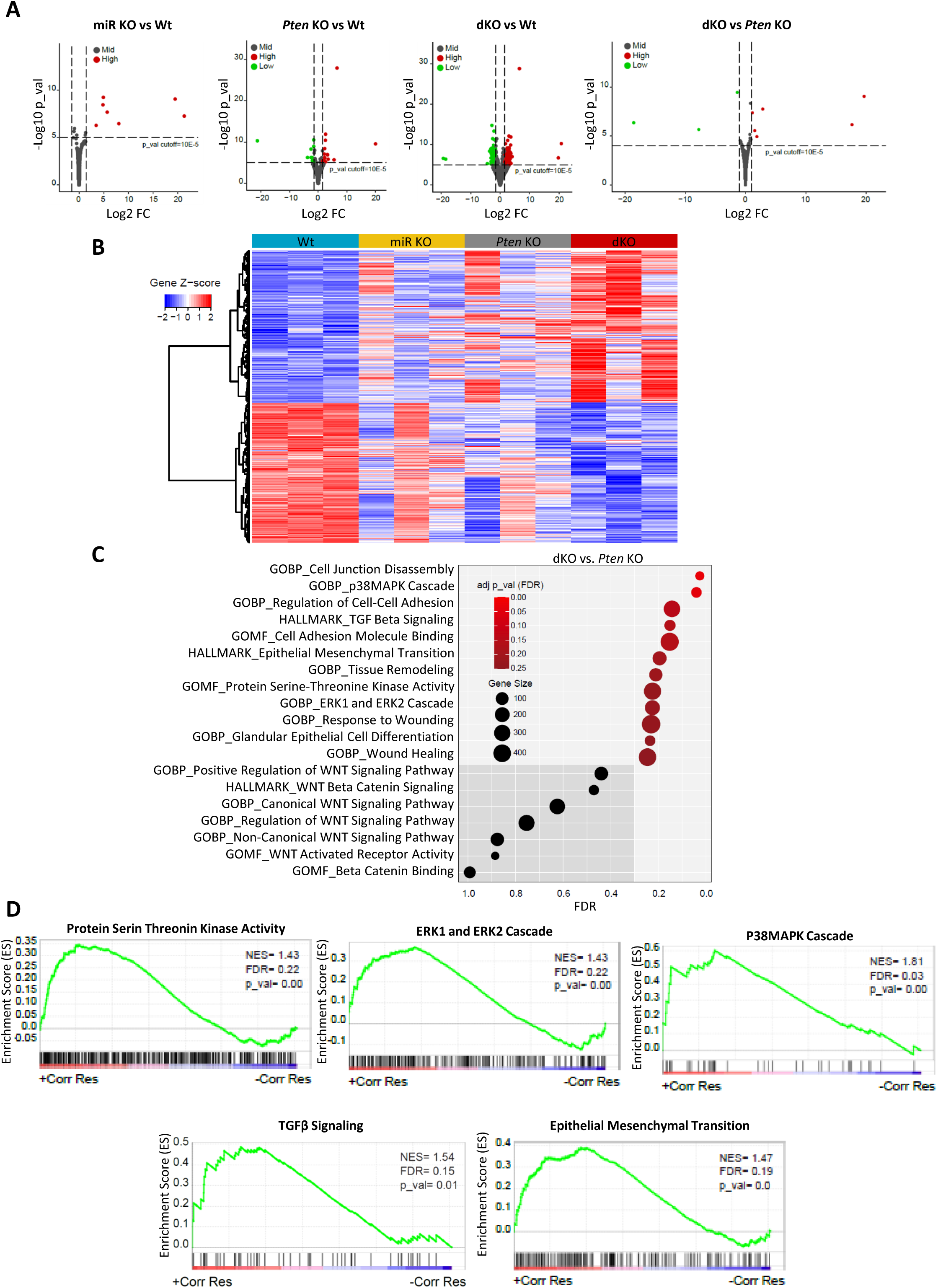
Transcriptome analysis of colon epithelium dedificent in Pten and/or miRNA miR-322/503. **(A)** Volcano plots of differentially expressed genes (DEGs) comparing Cre:ER(T)^-/-^ *Pten*^F/F^ miR-322/503^+/+^ (Wt), Cre:ER(T)^-/-^ *Pten*^F/F^ miR-322/503^-/-^(miR KO), Cre:ER(T)^+/-^ *Pten*^F/F^ miR-322/503^+/+^ (*Pten* KO) and Cre:ER(T)^+/-^ *Pten*^F/F^ miR-322/503^-/-^ (dKO) replicates, as indicated. Green dots represent significantly upregulated genes, while red dots indicate significantly downregulated genes. **(B)** Heatmap of hierarchical clustering analysis showing DEGs across Wt, *Pten* KO and dKO genotypes. **(C)** Dot plot representing gene set enrichment analysis (GSEA) of enriched gene signatures from the molecular signatures database (MSigDB) gene ontology (GO) biological processes (BP), molecular function (MF), cellular components (CC) and hallmarks comparing *Pten* KO and dKO genotypes. The plot highlights transcriptomic signatures deregulated in dKO colorectal tissue. Each red node represents a distinct positive enrichment, while blue nodes represent negative enrichment annotations; node size corresponds to the number of genes included within the annotation. **(D)** Enriched gene annotation from GSEA for “Protein Serin Threonin Kinase Activity”, “ERK1 and ERK2 Cascade”, “p38MAPK Cascade”, “Response to Transforming Growth Factor β” and “Epithelial Mesenchymal Transition”.

Given that the CRC development was dramatically increased in dKO over *Pten* KO colonic epithelium, we focused our analysis on comparing these two genotypes. Our GSEA primarily retrieved gene signatures associated with tissue remodeling processes such as Cell-cell Adhesion, Cell Adhesion Molecule Binding, Response to Wounding/Would healing or Epithelial-to-Mesenchymal Transition (EMT) **(Figure 4C-D)**. Regarding signaling pathway modifications, we found significative changes in the signatures associated with TGFβ signaling and the MAPK signaling pathways ERK1/2 and p38 (**Figure 4C-D)**. These pathways are also known to play critical roles in cellular growth and survival and are commonly dysregulated in CRC. These findings suggest that the loss of miR-322^∼^503 and *Pten* triggers a transcriptional program that actively promotes cell proliferation and EMT, likely through its impact on TGFβ and MAPK signaling.

### Colorectal tissue deficient in both Pten and miR-322^∼^503 exhibits hyperactivation and TGFβ signaling pathways and enhanced cyclinD1 expression

To analyze whether transcriptomic dysregulation of ERK1/2 and p38 MAPK signaling were translated to alterations in the activation of such pathways, we carried-out Western Blot analysis of protein lysates from colorectal epithelium from Wt, *miR* KO, *Pten* KO or dKO mice. The effect of *Pten* deletion on the PI3K/AKT pathway was investigated by analyzing the phosphorylated forms of AKT, p70S6K, and S6 as a read-out of pathway activation. As we mentioned before (**Supplementary Figure S1A)**, it is important to highlight that *Pten* deletion in epithelial cells occurs in a mosaic pattern, which explains the partial expression of PTEN observed in *Pten* KO and dKO conditions. Despite such mosaicism *Pten* deletion, a marked increase in the phosphorylated forms of AKT, p70S6K, and S6 was observed in all *Pten* KO and dKO conditions. Interestingly, dKO did not show further significant increase of AKT, p70S6K, and S6 phosphorylation over *Pten* KO conditions, suggesting that lack of miR-322^∼^503 does not affect PI3K/AKT activation directly **(Figure 5A)**. This result is further supported by the *miR* KO conditions, in which PI3K/Akt signaling pathway is not significantly increased over Wt ones. Regarding MAPK signaling activation, Western blot analysis revealed an elevated phosphorylation levels of JNK, p38, MKK4, and ERK1/2, in dKO conditions compared to the rest on genotypes, suggesting an activation of the three main MAPK signaling pathways **(Figure 5A, 5B)**. Among all three MAPKs, the ERK1/2 signaling pathway plays a pivotal role in development of CRC. Therefore, we further analyzed the increase in ERK1/2 phosphorylation by IHC staining of Wt, *miR* KO, *Pten* KO or dKO colon epithelium. Noteworthy, a marked increase of p-ERK1/2 staining was observed in lesions from dKO mice (**Figure 5B**). Moreover, analysis of cyclin D1 expression in consecutive tissue sections revealed a pronounced increase in lesions displaying increased ERK phosphorylation.

**Figure 5.**
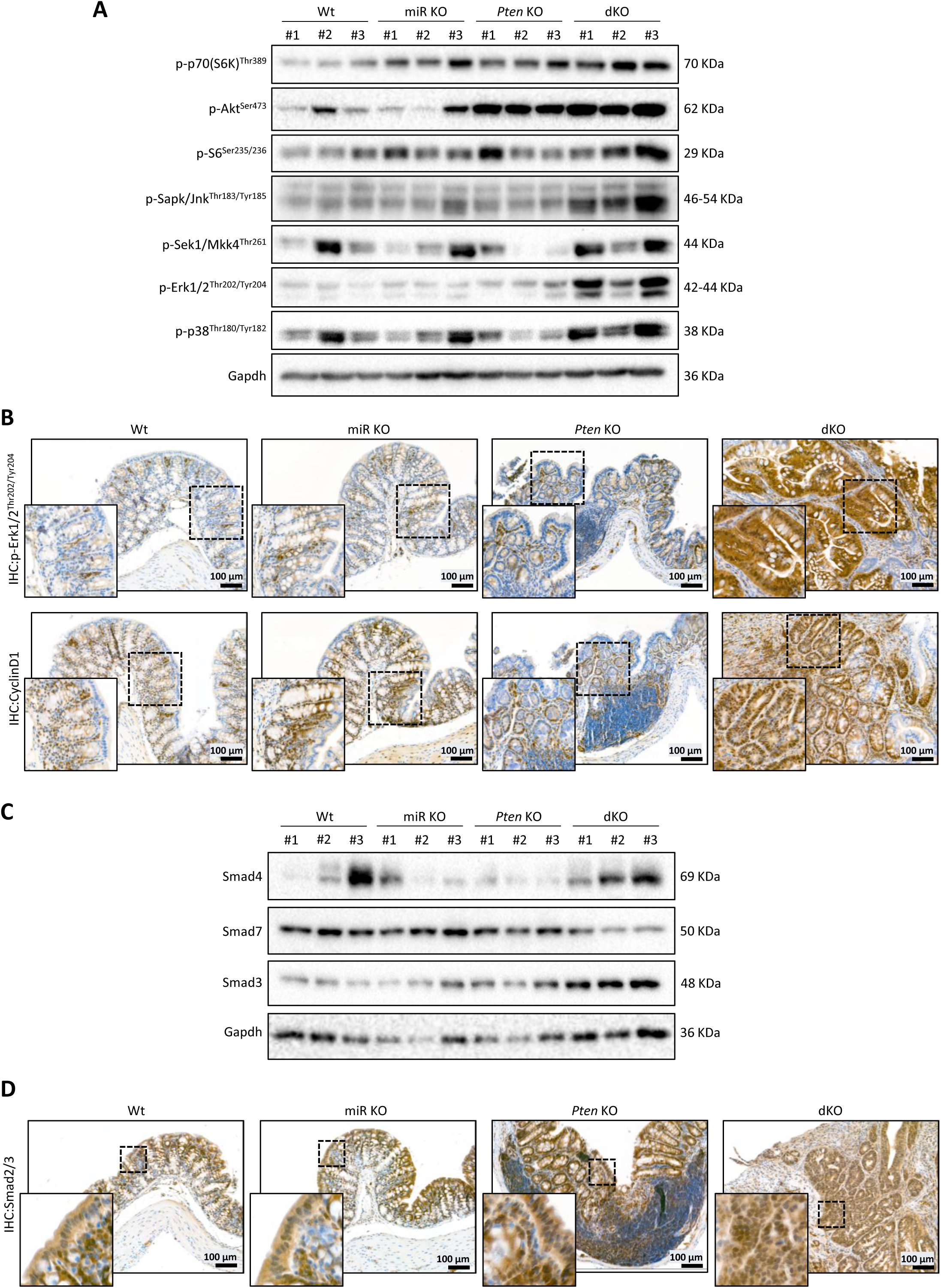
Double loss of *Pten* and miR-322/503 enhance MAPK and TGFβ signaling. **(A)** Western blot images of phosphorylated signaling proteins: p-p70(S6K)^Thr389^, p-Akt^Ser473^, p-S6^Ser235/236^, p-eif4e^Ser209^, p-Sapk/Jnk^Thr183/Tyr185^, p-Sek1/Mkk4^Thr261^, p-Erk1/2^Thr202/Tyr204^, p-p38^Thr180/Tyr182^ and Cyclin D1. The analysis was performed on Cre:ER(T)^-/-^ *Pten*^F/F^ miR-322/503^+/+^ (Wt), Cre:ER(T)^-/-^ *Pten*^F/F^ miR-322/503^-/-^ (miR KO), Cre:ER(T)^+/-^ *Pten*^F/F^ miR-322/503^+/+^ (*Pten* KO) and Cre:ER(T)^+/-^ *Pten*^F/F^ miR-322/503^-/-^(dKO) epithelial colorectal tissue. Membranes were re-probed with Gapdh as loading control. **(B)** Representative immunohistochemistry images of p-Erk1/2^Thr202/Tyr204^ and Cyclin D1 in colorectal sections from Wt, miR KO, *Pten* KO and dKO transgenic mice. Scale bar: 100μm (15X magnification). **(C)** Western blot images of showing the expression of SMAD3, SMAD4 and SMAD7. The analysis was performed on Wt, miR KO, *Pten* KO and dKO epithelial colorectal tissue. Membranes were re-probed with Gapdh as loading control. (D) Representative immunohistochemistry images of Smad2/3 in Wt, miR KO, *Pten* KO and dKO colorectal sections. Scale bar: 100μm (15X magnification).

Finally, we investigated alterations in signaling pathways activated in response to TGFβ. The SMAD family of transcription factors are one of the main transducers of cellular responses after TGFβ receptor engagement^28^. Among them, the SMAD3 and SMAD4 are the main transcription factors activated by TGFβ signaling, while SMAD7 is an inhibitory SMAD. Western blot analysis of Wt, *miR* KO, *Pten* KO or dKO derived lysates revealed global increase of SMAD3 and SMAD4 expression and a decrease in SMAD7 in dKO mice (**Figure 5C**). Moreover, immunohistochemistry showed an increased nuclear staining of SMAD2/3 in dKO tumors **(Figure 5D)**. This result suggests that TGFβ/SMAD signaling is enhanced by the lack of *Pten* and the miRNA miR-322^∼^503.

## DISCUSSION

MicroRNAs serve as key regulators of post-transcriptional gene expression and have been increasingly recognized as pivotal players in oncogenic processes^46^. While numerous miRNAs have been associated with CRC pathogenesis, the functional characterization of individual miRNAs – particularly their *in vivo* roles – remains an underexplored area of research. This study addresses a critical knowledge gap by investigating the miR-424(322)^∼^503 cluster through a genetically engineered mouse model, revealing its essential role in counteracting PTEN deficiency-driven CRC progression. Our findings demonstrate that genetic ablation of miR-424(322)^∼^503 markedly accelerates colorectal tumorigenesis in *Pten*-deficient mice, establishing this miRNA cluster as a crucial tumor suppressor in the *Pten*-loss context. The tumor-suppressive mechanism appears to involve coordinated regulation of two fundamental signaling axes: the ERK-MAPK signaling, a pathway frequently hyperactivated in human CRCs and; the TGFβ/Smad signaling, a critical pathway maintaining epithelial differentiation and apoptotic responses. Notably, the combined loss of *Pten* and miR-424(322)^∼^503 creates a permissive environment for malignant transformation by simultaneously disrupting these interconnected networks. This dual pathway dysregulation likely explains the observed acceleration of adenoma-to-adenocarcinoma progression in our model, providing mechanistic insights into how miRNA-mediated regulation protects intestinal epithelium integrity under oncogenic stress conditions.

One of the most intriguing questions about miR-424(322)^∼^503 is its paradoxical role as oncogene or tumour suppressor gene depending on the cellular context^16–18^. However, tumor suppressive and oncogenic functions have been demonstrated in independent models, leaving the possibility that different experimental scenarios may affect the function of miR-424(322)^∼^503. We have recently demonstrated that miR-424(322)^∼^503 acts as oncogene in EC initiated by *Pten* deletion by regulating cell proliferation and apoptosis. Here, using the same mouse model, we have found that miR-424(322)^∼^503 functions as tumor suppressor in CRC initiated by *Pten* deficiency. These completely opposite *in vivo* effects in the endometrium and the colonic epithelium highlights the duality of miR-424(322)^∼^503 depending on the tissue, even when carcinogenesis is initiated by the same driver. Furthermore, the function of this miRNA specifically in CRC is still controversial and both oncogenic ^20,21^ and tumor suppressive functions has been proposed^22–24^. Another result that deserves discussion is the effect of *Pten* ablation on miR-424(322)^∼^503 expression in colorectal epithelial cells. Our previous results demonstrated that loss of *Pten* or TGFβ caused an up-regulation of miR-424(322)^∼^503 expression, which acts as an oncogene by promoting EC progression^25^. Similarly, we found that *Pten* or TGFβ increase the expression of miR-424(322)^∼^503, which ultimately has the contrary function to that found in the endometrium. This result strengthens the importance of PI3K/Akt and TGFβ signaling pathways as critical regulators of miR-424(322)^∼^503 expression, independently of its function in the target tissue.

At the heart of CRC pathogenesis remain disrupted signaling pathways that promote tumor formation, support cancer cell growth, and facilitate metastatic spread. Among them, the Wnt/β-catenin, the RAS/RAF/ERK, the PI3K/AKT, and the TGF-β are frequently altered by a combination of genetic mutations^29^. It has been recently demonstrated that miR-424(322)^∼^503 cluster can modulate β-catenin signnalling by targeting LRP6 receptor in mammary epithelium ^26^. Based these findings and the crucial role of β-catenin signaling in CRC^30^, it was easy to speculate that CRC caused by double Pten and miR-424(322)^∼^50 ablation was caused by dysregulation of classic β-catenin signaling. However, no changes in β-catenin expression of cellular localization nor in LRP6 expression have been found in double *Pten* and miR-424(322)^∼^503 KO mice, suggesting this pathway is not participating in our model. Instead, our findings demonstrate that miR-424(322)^∼^503 knockout enhances MAPK signaling in Pten deficient colorectal cells, resulting in an epithelium with exacerbated activation of both PI3K/Akt and MAPK pathways. Deregulation of the Ras/Raf/MEK/ERK MAPK signaling pathway is a critical factor that drives the progression of CRC^31^. This pathway acts as a critical signaling hub that governs cell proliferation and cell cycle progression. Among human cancers, somatic *RAS* mutations occur in roughly 30% of cases, triggering constitutive activation of the downstream kinase cascade. The process begins with mutant RAS activating RAF, which phosphorylates MEK, ultimately leading to sustained MAPK/ERK signaling. This stepwise amplification drives uncontrolled cellular growth and survival. The cooperation between the PI3K/Akt and MAPK signaling pathways is well-documented in CRC, as both pathways play critical roles in tumor initiation, progression, metastasis and resistance to therapy^29^. It is worth mentioning that besides RAS/RAF/ERK MAPK signaling, we also have found increased phosphorylation of p38^32^ and JNK^33,34^, two other MAPK that can participate CRC development and progression. These findings collectively underscore the critical role of PI3K/Akt and MAPK pathway cooperation in CRC pathogenesis and highlight their potential as therapeutic targets.

Finally, we have identified signatures associated with TGFβ signaling EMT, two closely interconnected processes involved in carcinogenesis^35^. Regarding TGFβ signaling, the increased expression of SMAD3 and SMAD4, along with the decreased expression of SMAD7 and the enhanced nuclear accumulation of SMAD2/3 in colorectal cancer (CRC) derived from miRNA and *Pten* double knockout mice, strongly suggests an activation of the pathway. The upregulation of TGFβ signaling in tumors is closely linked to EMT, as these two processes are highly interconnected and play critical roles in cancer progression. Therefore, it is reasonable to speculate that the increased EMT signatures are associated with the activation of TGFβ/SMAD signaling.

In conclusion, this study emphasizes the complex and context-dependent role of the miR-424(322)∼503 cluster in CRC, highlighting its critical involvement in regulating key signaling pathways such as MAPK and TGFβ/SMAD. By demonstrating how the loss of this miRNA cluster accelerates tumorigenesis in a Pten-deficient mouse model, we provide important mechanistic insights into its tumor-suppressive function in CRC. Furthermore, our findings underscore the intricate interplay between PI3K/Akt, MAPK, and TGFβ signaling pathways in driving cancer progression, revealing potential therapeutic targets for the treatment of CRC. The paradoxical dual role of miR-424(322)∼503, acting as both an oncogene and tumor suppressor depending on the tissue context, opens new avenues for future research aimed at understanding its precise molecular mechanisms.

## MATERIAL AND METHODS

### Experimental Mouse Models

Mice were housed in a barrier facility and pathogen-free procedures were used in all mouse rooms. Animals were under 12 hours of light/dark cycles at 22°C, and they had *ad libitum* access to water and food. All procedures were performed according to the guidelines of the Ethical Committee of Universitat de Lleida and the National Institute of Health Guide for the Care and Use of Laboratory Animals. Conditional *Pten* knockout (C; 129S4-Ptentm1Hwu/J or *Pten*^F/F^) and Cre:ER(T) mice were obtained from the Jackson Laboratory (Bar Harbor, ME, USA). miR-322/503^-/-^ (FVB/NJ) mice were a gift from Prof. Jose Silva. Cre:ER(T) *Pten*^F/F^ miR-322/503^-/-^ mice were generated by crossing Cre:ER(T)^+/-^ *Pten*^F/F^ and miR-322/503^-/-^ mice. Three weeks after birth, mice were weaned and genotyped as previously described ^23^^36 38^. Genotyping primers and PCR conditions are available in Supplementary Materials (Table S1).

### Tamoxifen Administration

Tamoxifen (T5648, Sigma-Aldrich) was prepared and administered as previously described^36^. Briefly, tamoxifen powder was dissolved in 100% ethanol at a concentration of 100mg/ml. Then, tamoxifen was emulsified in corn oil (C8267, Sigma-Aldrich) at a final concentration of 10mg/ml. Mice between 5 and 8 weeks were injected with a single 0.5mg of tamoxifen intraperitoneally.

### Isolation of Epithelial Colon Cells and Organoid Culture

Mice were euthanized by cervical dislocation and the colorectal portions compressed between cecum and rectum were dissected. Colon portions were longitudinally opened and gently washed in ice-cold Phosphate Buffered Saline (PBS). Colons were incubated in digestion medium (50mg/ml collagenase type II (17101-015, Gibco) and 10% Fetal Bovine Serum (FBS) (A52567-01, Gibco) in DMEM (41965-039, Gibco)) for 2 hours at 37°C. To isolate colon crypts, digested colons were passed through a 70μm sterile strainer and washed in DMEM/F12 (11039-021, Gibco). When *Pten* ablation was required, colon crypts were incubated for 30 minutes at 37°C with TAT-Cre peptide ^36^ and gently washed in DMEM/F12 before seeding. Colon crypts were resuspended in IntestiCult™ Organoid Growth Medium (Mouse) (06005, STEMCELL) containing 50% Matrigel™ (354234, Corning) to obtain 1500 crypts/ml, and plated as domes in a multiwell plate. Domes containing 50% Matrigel™ were polymerized for 10 minutes at 37°C and then, IntestiCult™ Organoid Growth Medium (Mouse) was added to cover the dome. Every 2-3 days medium was replaced until organoids were completely formed. Organoid treatments were performed two days after medium replacement. TGFβ (#PHG9214, Gibco) treatments were performed at 10ng/μl for 16 hours.

### miRNA Extraction and Real-Time qPCR

miRNA extraction was performed in total epithelial colon tissue, isolated polyps and colon organoids using mirVana miRNA isolation kit (AM1561, Ambion) following manufacturer’s instructions. miRNA extracts were quantified with NanoPhotometer (N60 UV/Vis Spectrophotometer, IMPLEN) and stored frozen at -80°C. Reverse transcription was performed using 50ng of RNA using High-Capacity cDNA Reverse Transcription (4368815, Applied Biosystems), according to manufacturer’s protocol. Retrotrancription was performed at 16°C for 30 minutes, followed by 60 cycles of 20°C for 3 seconds, 42°C for 3 seconds and 50°C for 2 seconds, reaction was inactivated at 85°C for 5 minutes. Primers for retrotranscription are detailed in Supplementary Materials (Table S2). Quantitative real-time PCR detection of miRNAs was performed with the CFX96 (BioRad) using PowerUp SYBR Green MasterMix (A25742, Applied Biosystems). Real-time PCR was performed at 95°C for 3 minutes, followed by 40 cycles of 95°C for 10 seconds and 60°C for 30 seconds, and finished with a descendent gradient of temperature. Sequences of primers used in real-time PCR are shown in Supplementary Materials (Table S3). Relative expressions were determined from cycle threshold (Ct) values, which were normalized to sno-202 as a housekeeping miRNA. Experiments were performed at least three times, and every group was performed in triplicate.

### Total mRNA Extraction and Real-Time qPCR

Total RNA from colon epithelial tissue was extracted using SurePrep TrueTotal RNA Purification kit (BP2800-50, Fisher BioReagents) following manufacturer’s instructions. Total RNA extracts were quantified with NanoPhotometer and stored frozen at -80°C. Quantitative real-time PCR was performed with 50ng of total RNA using the one-step protocol qPCRBIO Probe 1-step Go (PB25.44-01, PCR Biosystems) according to manufacturer’s protocol. Primers used for gene expression assays were commercially obtained from Applied Biosystems and are listed in Supplementary Materials (Table S4). Relative expressions were assessed by cycle threshold (Ct) values, which were normalized to *Gapdh* expression as an internal control. Experiments were performed at least three times, and every group was performed in triplicate.

### Whole Genome mRNA Sequencing

2×150bp mRNA sequencing was performed at GeneWiz (Azenta Life Sciences) on NovaSeq equipment at a depth of 20M reads per sample. Quality control of all FASTQ files was performed using the FASTQC tool (v0.11.9) (reference) and MultiQC tool implemented in R. For mRNAseq quantification the mapping-based mode of Salomon (v1.5.2) (reference) was used to perform transcript-level quantification with default parameters, using the tximport library (v1.28.0) (reference). Low-expression genes, defined as those with fewer than 10 reads across all samples, were filtered out to prior to analysis. Data normalization and differential expression analysis between miR KO, *Pten* KO, dKO and Wt conditions were performed using the DESeq2 R package (v1.30.1) (reference). To conduct Gene Set Enrichment Analysis (GSEA reference), genes from the dKO vs Wt comparison were ranked by foldchange and analyzed using the msigdbr R function. Selected pathways were visualized in a dot plot, displaying FDR values and gene set sizes, created using ggplot2 package in R.

### Western blot

Western blot analysis was performed as previously described ^36^ with minor variations. To isolate total colorectal epithelium, colons were dissected, opened longitudinally and washed twice with PBS. Epithelial cells were isolated with a scalpel and lysed with 2% SDS, 125mM Tris-HCl pH 6.8. Relative protein concentrations were determined with a colorimetric protein assay kit (5000112, BioRad). Equal amounts of protein were loaded to an acrylamide gel and transferred to PDVF membranes (IPVH00010, Millipore). To avoid nonspecific antibody binding, membranes were blocked for 1 hour with 5 % non-fat milk in TBS-T (20mM Tris-HCl pH7.4, 150 mM NaCl, 0.1% Tween-20). Membranes were incubated with primary antibodies during 16 hours at 4°C. Then, membranes were incubated for 1 hour with secondary antibodies at room temperature. Finally, signal was detected with Immobilon Forte Western HRP Substrate (WBLUF0100, Millipore). Primary and secondary antibodies used for Western blot and their concentrations are detailed in Supplementary Materials (Table S5). Intensity band quantification was performed in an image analyzer (ImageJ, version 1.46r; NIH, Bethesda, MD; USA).

### Tissue Processing and Immunohistochemistry Analysis on Paraffin Sections

Mice were euthanized and colons were collected, formalin-fixed overnight at 4°C and paraffin-embedded. Then, paraffin sections of 3 μm were dried for an hour at 80°C, dewaxed in xylene, gradually rehydrated in ethanol and washed in PBS. For antigen retrieval, slides were incubated in EnVision FLEX and high pH or low pH solutions (K8004 orK8005, DAKO) for 20 minutes at 95°C, depending on each antibody. Then, slides were blocked with endogenous peroxidase through 3% H_2_O_2_ incubation and washed three times in PBS. Then, primary antibodies were incubated for 30 minutes at room temperature, washed in PBS and incubated with Horseradish Peroxidase (HRP)-conjugated secondary antibodies. Staining was visualized through reaction with EnVision Detection Kit (K4065, DAKO) using diaminobenzidine (DAB) substrate. Finally, slides were counterstained with Harrys hematoxylin. All primary and secondary antibodies used for immunohistochemistry and their concentrations are listed in Supplementary Materials (Table S6).

### Statistical Analysis

Statistical analysis was performed according to each experiment. Shortly, comparisons between two groups were analyzed with Student’s *t*-test and represented as mean ± standard error for the mean. Contingency tables were analyzed by χ^2^-test, followed by Fisher’s exact test. All experiments were performed at least three times, and all experimental groups were done in triplicates. For statistical analysis, GraphPad Prism (Version 8.0, Graphpad Software, Inc.) was used.

## Supporting information

Supplementary FMethods

## ACKNOWLEDGEMENTS

This study has been funded by the Institut de Recerca Biomédica de Lleida (PIRS2023, XD), Ministerio de Ciencia, Innovación y Universidades (PID2022-141220OB-I00, XD and PID2019-104734RB-I00, XD), and Instituto de Salud Carlos III (ISCIII) (PI21/00672, DL-N and PI24/01255, DL-N) (co-funded by the European Regional Development Fund. ERDF, a way to build Europe), and by the CIBERonc network (XM-G, CB16/1200231). We thank the Generalitat of Catalonia, Agency for Management of University and Research Grants (2021SGR01609 and 2021SGR01098). The authors also want to thank the CERCA programme / Generalitat de Catalunya for institutional support.

## AUTHOR CONTRIBUTIONS

MV-S, DL-N, and XD designed the research and developed the project concept. MV-S, DL-N, and XD prepared the manuscript. JE, ME, RR-B, JMS, XM-G, DL-N, and XD provided essential materials and resources necessary for conducting the research. MV-S,

NB, RN, JT and XD performed the experiments and collected the data. MV-S, NB, RN, DL-N, and XD analyzed the data and conducted statistical analyses. All authors contributed to data interpretation and critically revised the manuscript.

## FIGURE LEGENDS

**Figure S1.**
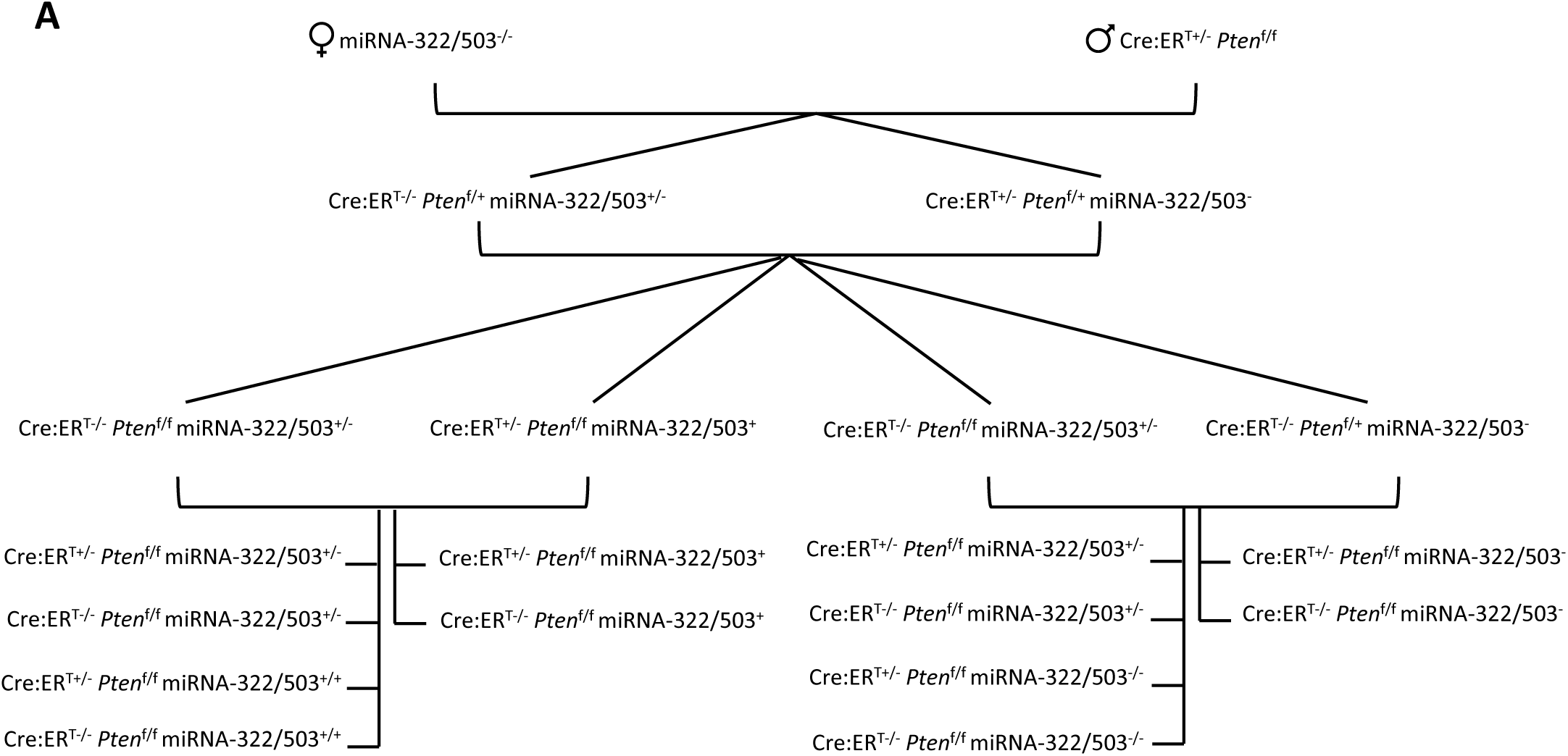

**Figure S2.**
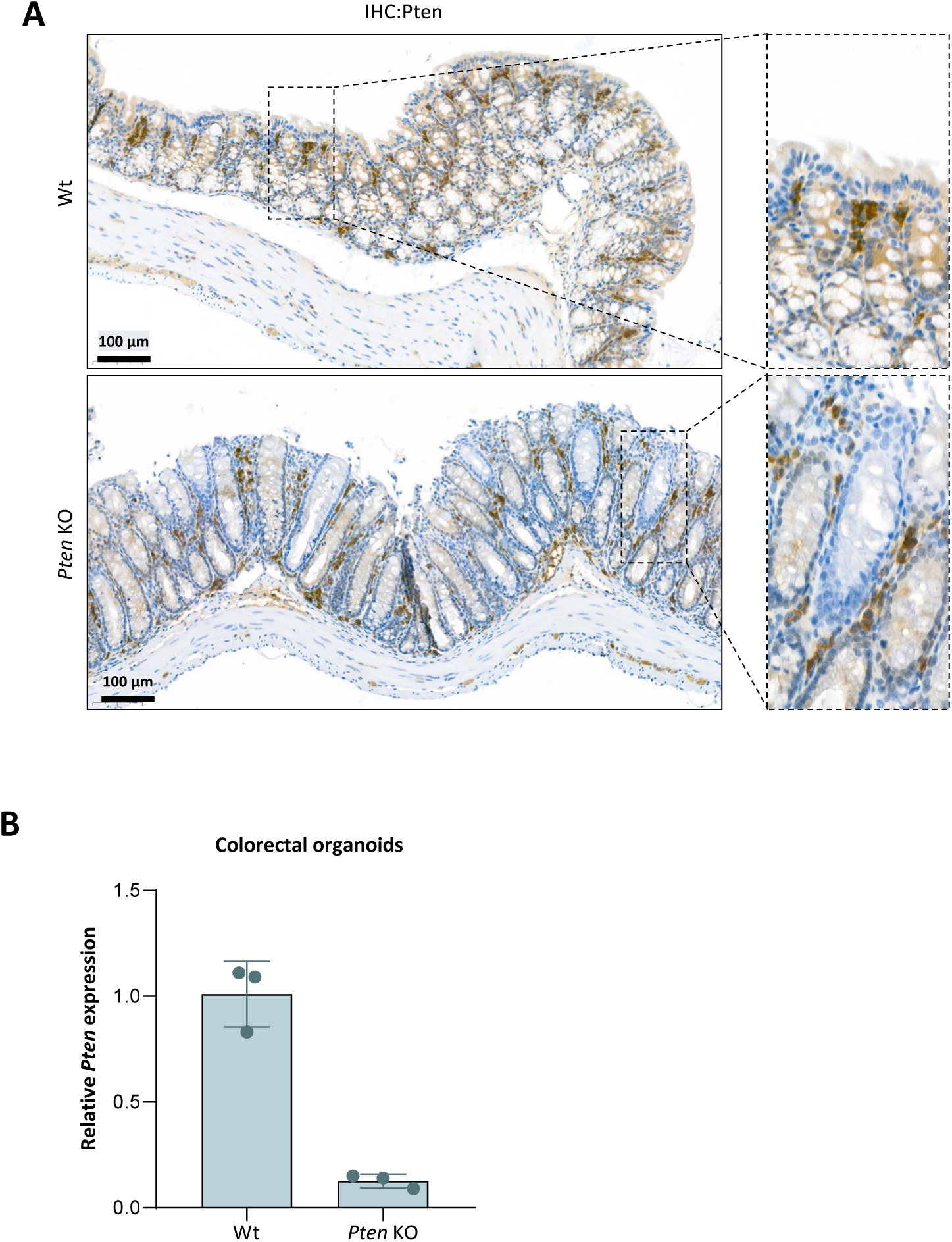

## REFERENCES.

1. Bray, F., et al. Global cancer statistics 2022: GLOBOCAN estimates of incidence and mortality worldwide for 36 cancers in 185 countries. CA: A Cancer Journal for Clinicians 74, 229–263 (2024).

2. GLOBACAN 2022. https://gco.iarc.who.int/today/.

3. Ahadi, M., Sokolova, A., Brown, I., Chou, A. & Gill, A. J. The 2019 World Health Organization Classification of appendiceal, colorectal and anal canal tumours: an update and critical assessment. Pathology 53, 454–461 (2021).

4. Nagtegaal, I. D. et al. The 2019 WHO classification of tumours of the digestive system. Histopathology 76, 182–188 (2020).

5. Dekker, E., Tanis, P. J., Vleugels, J. L. A., Kasi, P. M. & Wallace, M. B. Colorectal cancer. The Lancet 394, 1467–1480 (2019).

6. Guinney, J., et al. The consensus molecular subtypes of colorectal cancer. Nat Med 21, 1350–1356 (2015).

7. Rejali, L., et al. Principles of Molecular Utility for CMS Classification in Colorectal Cancer Management. Cancers 15, 2746 (2023).

8. Li, J., Ma, X., Chakravarti, D., Shalapour, S. & DePinho, R. A. Genetic and biological hallmarks of colorectal cancer. Genes & Development 35, 787 (2021).

9. Nguyen, L. H., Goel, A. & Chung, D. C. Pathways of Colorectal Carcinogenesis. Gastroenterology 158, 291 (2019).

10. Ross, M. H. & Pawlina, Wojciech. Histología Texto y Atlas Color Con Biología Celular y Molecular. (Médica Panamericana, Buenos Aires, 2013).

11. Serebriiskii, I. G., et al. Comprehensive characterization of PTEN mutational profile in a series of 34,129 colorectal cancers. Nature Communications 13, 1618 (2022).

12. Zhou, X.-P. et al. PTEN Mutational Spectra, Expression Levels, and Subcellular Localization in Microsatellite Stable and Unstable Colorectal Cancers. Am J Pathol 161, 439–447 (2002).

13. Li, X.-H., et al. PTEN expression and mutation in colorectal carcinomas. Oncol Rep 22, 757–764 (2009).

14. Day, F. L., et al. *PIK3CA* and *PTEN* Gene and Exon Mutation-Specific Clinicopathologic and Molecular Associations in Colorectal Cancer. Clinical Cancer Research 19, 3285–3296 (2013).

15. Liu, Q., et al. miR-16 family induces cell cycle arrest by regulating multiple cell cycle genes. Nucleic Acids Res 36, 5391–5404 (2008).

16. Ghafouri-Fard, S., Askari, A., Hussen, B. M., Taheri, M. & Akbari Dilmaghani, N. Role of miR-424 in the carcinogenesis. Clin Transl Oncol 26, 16–38 (2024).

17. Li, S., Wu, Y., Zhang, J., Sun, H. & Wang, X. Role of miRNA-424 in Cancers. Onco Targets Ther 13, 9611–9622 (2020).

18. Wu, Y., Wang, W., Yang, A.-G. & Zhang, R. The microRNA-424/503 cluster: A master regulator of tumorigenesis and tumor progression with paradoxical roles in cancer. Cancer Lett 494, 58–72 (2020).

19. Wang, F., et al. H19X-encoded miR-424(322)/-503 cluster: emerging roles in cell differentiation, proliferation, plasticity and metabolism. Cell Mol Life Sci 76, 903–920 (2019).

20. Li, N. CircTBL1XR1/miR-424 axis regulates Smad7 to promote the proliferation and metastasis of colorectal cancer. J Gastrointest Oncol 11, 918–931 (2020).

21. Dai, W., et al. miR-424-5p promotes the proliferation and metastasis of colorectal cancer by directly targeting SCN4B. Pathol Res Pract 216, 152731 (2020).

22. Oneyama, C., et al. MiR-424/503-mediated Rictor upregulation promotes tumor progression. PLoS One 8, e80300 (2013).

23. Ghonbalani, Z. N., Shahmohamadnejad, S., Pasalar, P. & Khalili, E. Hypermethylated miR-424 in Colorectal Cancer Subsequently Upregulates VEGF. J Gastrointest Cancer 53, 380–386 (2022).

24. Liu, Y., et al. miR-424-5p reduces 5-fluorouracil resistance possibly by inhibiting Src/focal adhesion kinase signalling-mediated epithelial-mesenchymal transition in colon cancer cells. J Pharm Pharmacol 73, 1062–1070 (2021).

25. Vidal-Sabanes, M., et al. Endometrial cancer progression driven by PTEN-deficiency requires miR-424(322)∼503. 2025.04.07.647575 Preprint at 10.1101/2025.04.07.647575 (2025).

26. Nekritz, E. A., et al. miR-424/503 modulates Wnt/β-catenin signaling in the mammary epithelium by targeting LRP6. EMBO Rep 22, e53201 (2021).

27. Zhao, H., et al. Wnt signaling in colorectal cancer: pathogenic role and therapeutic target. Mol Cancer 21, 144 (2022).

28. Aashaq, S., et al. TGF-β signaling: A recap of SMAD-independent and SMAD-dependent pathways. J Cell Physiol 237, 59–85 (2022).

29. Li, Q., et al. Signaling pathways involved in colorectal cancer: pathogenesis and targeted therapy. Sig Transduct Target Ther 9, 1–48 (2024).

30. Fearon, E. R. & Vogelstein, B. A genetic model for colorectal tumorigenesis. Cell 61, 759–767 (1990).

31. Fang, J. Y. & Richardson, B. C. The MAPK signalling pathways and colorectal cancer. Lancet Oncol 6, 322–327 (2005).

32. Grossi, V., Peserico, A., Tezil, T. & Simone, C. p38α MAPK pathway: a key factor in colorectal cancer therapy and chemoresistance. World J Gastroenterol 20, 9744–9758 (2014).

33. Lee, Y.-H., et al. JNK-mediated Ser27 phosphorylation and stabilization of SIRT1 promote growth and progression of colon cancer through deacetylation-dependent activation of Snail. Mol Oncol 16, 1555–1571 (2022).

34. Fang, M., et al. IL33 Promotes Colon Cancer Cell Stemness via JNK Activation and Macrophage Recruitment. Cancer Res 77, 2735–2745 (2017).

35. Lee, J. H. & Massagué, J. TGF-β in developmental and fibrogenic EMTs. Semin Cancer Biol 86, 136–145 (2022).

36. Navaridas, R., et al. Transient and DNA-free in vivo CRISPR/Cas9 genome editing for flexible modeling of endometrial carcinogenesis. Cancer Commun (Lond) (2023) doi:10.1002/cac2.12409.

