## Supplementary figures and images for "miR-424(322)^∼^503 impairs colon cancer progression driven by PTEN deficiency"

### Supplementary Figures

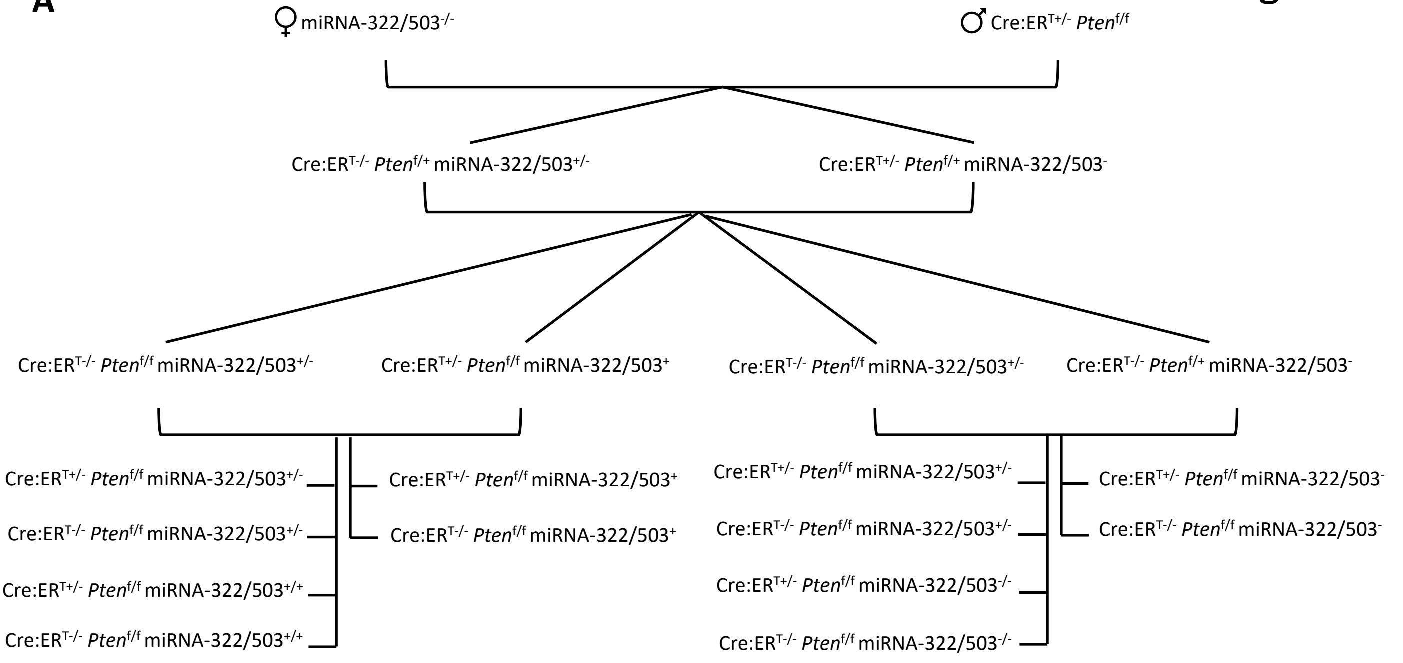

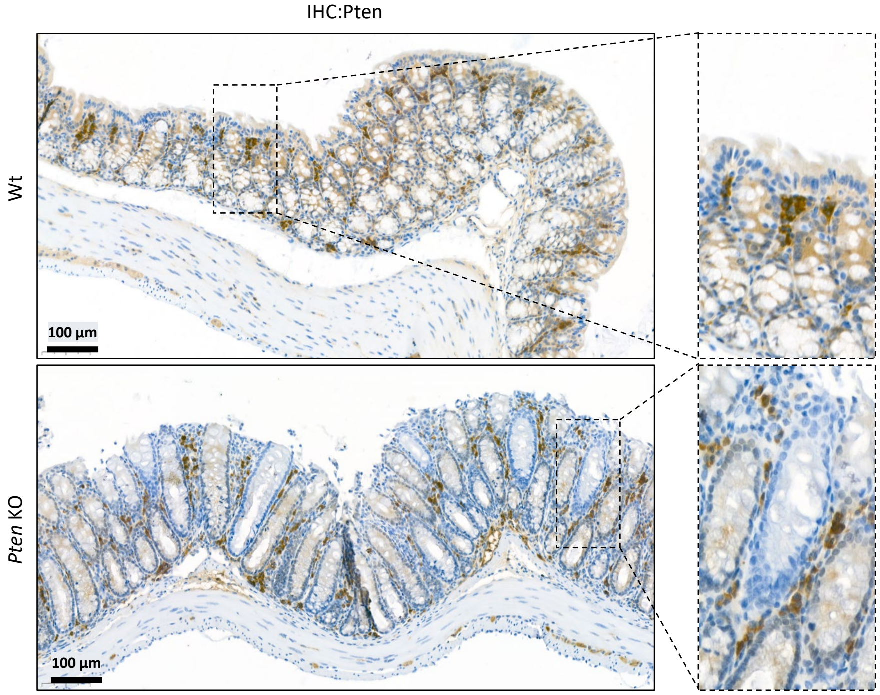

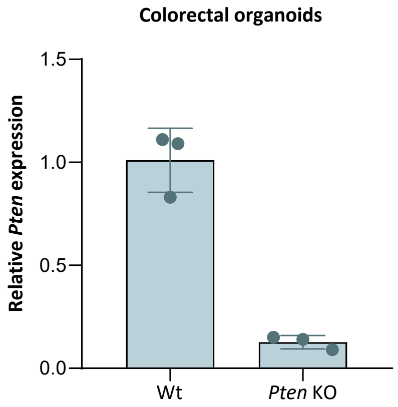
