## Supplementary FMethods for "miR-424(322)^∼^503 impairs colon cancer progression driven by PTEN deficiency"

| **Allele** | **Primers** | | **PCR conditions** | | | **Product** | |
| --- | --- | --- | --- | --- | --- | --- | --- |
|  |  |  | **Temperature** | **Time** | **Cycles** | **Genotype** | **Weight** |
| **Cre:ER^T^** | Fwd: | ACGAACCTGGTCGAAATCGTGCG | 94 ºC | 2 min | 1 | Cre:ER^T-/-^ | No band |
|  |  |  | 94 ºC | 45 sec | 32 |  |  |
|  |  |  | 65 ºC | 45 sec |  |  |  |
|  | Rev: | CGGTCGATGCAACGAGTGATGAG | 72 ºC | 45 sec |  | Cre:ER^T+/-^ | 350 bp |
|  |  |  | 72 ºC | 5 min | 1 |  |  |
|  |  |  | 4 ºC | ∞ |  |  |  |
| ***Pten*^f/f^** | Fwd: | CAAGCACTCTGCGAACTGAG | 94 ºC | 3 min | 1 | *Pten*^+/+^ | 156 bp |
|  |  |  | 94 ºC | 30 sec | 35 |  |  |
|  |  |  | 60 ºC | 1 min |  | *Pten*^f/+^ | 156 bp & 328 bp |
|  | Rev: | AAGTTTTTGAAGGCAAGATGC | 72 ºC | 2 min |  |  |  |
|  |  |  | 72 ºC | 2 min | 1 | *Pten*^f/f^ | 328 bp |
|  |  |  | 4 ºC | ∞ |  |  |  |
|  |  |  | 4 ºC | ∞ |  |  |  |
| **miR-322/503^+/+^** | Fwd: | CACCAGCAGATCCTGGAAAT | 94 ºC | 2 min | 1 | miR^+/+^ | 500 bp |
|  |  |  | 94 ºC | 30 sec | 35 |  |  |
|  |  |  | 59 ºC | 30 sec |  |  |  |
|  | Rev: | CAAGTGAGGCGCTAACAACA | 72 ºC | 2 min |  | miR^-/-^ | No band |
|  |  |  | 72 ºC | 5 min | 1 |  |  |
|  |  |  | 4 ºC | ∞ |  |  |  |
| **miR-322/503^-/-^** | Fwd: | AGTTTCGAAAAACGGACATA | 94 ºC | 2 min | 1 | miR^+/+^ | No band |
|  |  |  | 94 ºC | 30 sec | 35 |  |  |
|  |  |  | 59 ºC | 30 sec |  |  |  |
|  | Rev: | ATTGCCTCTCATTGTACCAC | 72 ºC | 2 min |  | miR^-/-^ | 700 bp |
|  |  |  | 72 ºC | 5 min | 1 |  |  |
|  |  |  | 4 ºC | ∞ |  |  |  |

**Table S1**. Genotyping primers and PCR conditions.

**Table S2.** Primers for reverse transcription of miR-322, miR-503 and sno-202.

| **Primer** | **Sequence** |
| --- | --- |
| Fwd Mmu-miR-322 | ACACTCCAGCTGGGCAGCAGCAATTCATGT |
| Fwd Mmu-miR-503 | ACACTCCAGCTGGGCAGCAGCAATTCATGT |
| Fwd sno-202 | ACACTCCAGCTGGGGCTGTACTGACTT |
| Common Reverse | GTGTCGTGGAGTCGGC |

**Table S3.** Primers for real-time PCR of miR-322, miR-503 and sno-202.

| **Primer** | **Sequence** |
| --- | --- |
| miR-322 | ACACTCCAGCTGGGCAGCAGCAATTCATGT |
| miR-503 | ACACTCCAGCTGGGCAGCAGCAATTCATGT |
| sno-202 | ACACTCCAGCTGGGGCTGTACTGACTT |

**Table S4**. Primers used for gene expression assays.

| **Gene** | **Company** | **Catalogue Number** |
| --- | --- | --- |
| ***Cdh2*** | Applied Biosystems | Mm01162497_m1 |
| ***Ctnnb1*** | Applied Biosystems | Mm00483039_m1 |
| ***Id1*** | Applied Biosystems | Mm00775936_g1 |
| ***Lrp6*** | Applied Biosystems | Mm00999795_m1 |
| ***Smad4*** | Applied Biosystems | Mm03023996_m1 |
| ***Smad7*** | Applied Biosystems | Mm00484742_m1 |
| ***Tgfb1*** | Applied Biosystems | Mm001178820_m1 |
| ***Vim*** | Applied Biosystems | Mm01333430_m1 |
| ***Gapdh*** | Applied Biosystems | Mm99999915_g1 |

**Table S5.** Antibodies used for Western blot analysis.

| **Antibody** | **Dilution** | **Catalogue Number, Company** |
| --- | --- | --- |
| α-p-Akt^Ser473^ | 1:1000 | 9271, Cell Signaling Technology |
| α-p-S6^Ser235/236^ | 1:1000 | 2211, Cell Signaling Technology |
| α-p-p70(S6K)^Thr389^ | 1:1000 | 9205, Cell Signaling Technology |
| α-p-Sapk/Jnk^Thr183/Tyr185^ | 1:1000 | 9251, Cell Signaling Technology |
| α-p-Sek1/Mkk4^Thr261^ | 1:1000 | 9156, Cell Signaling Technology |
| α-p-Erk1/2^Thr202/Tyr204^ | 1:1000 | 675502, Biolegend |
| α-p-p38^Thr180/Tyr182^ | 1:1000 | 903501, Biolegend |
| α-Smad3 | 1:1000 | ab28379, Abcam |
| α-Smad4 | 1:1000 | sc-7966, Santa Cruz |
| α-Smad7 | 1:1000 | 25840-1-AP, Proteintech |
| α-Gapdh | 1:10000 | ab8245, Abcam |
| α-IgG mouse-HRP | 1:10000 | 115-035-003 Jackson |
| α-IgG rabbit-HRP | 1:10000 | 111-035-003, Jackson |

**Table S6.** Antibodies used for immunohistochemistry assays.

| **Antibody** | **Dilution** | **Catalogue Number, Company** | **Antigen Retrieval** | **Secondary Antibody** |
| --- | --- | --- | --- | --- |
| α-β-catenin | RTU | IR702, Dako | High pH | EV FLEX Kit |
| α-Cyclin D1 | 1:25 | M3642, Dako | High pH | EV FLEX Kit |
| α-Lrp6 | 1:100 | 3395, Cell Signaling Technology | High pH | α-rabbit-biotin |
| α-N-cadherin | 1:250 | 22018-1-AP, Proteintech | High pH | α-rabbit-biotin |
| α-p-Erk^Thr202/Tyr204^ | 1:200 | 675502, Biolegend | High pH | α-mouse-biotin |
| α-Pten | 1:100 | M3627, Dako | High pH | EV FLEX Kit |
| α-Smad2/3 | 1:100 | sc-133098, Santa Cruz | High pH | α-mouse-biotin |
| α-IgG rabbit-biotin | 1:200 | 111-065-144, Jackson |  |  |
| α-IgG mouse-biotin | 1:200 | ab97363, Abcam |  |  |
| Streptavidin-HRP | 1:400 | P0397, Dako |  |  |
